# Divergence in the face of gene flow in two *Charadrius* plovers along the Chinese coast

**DOI:** 10.1101/406041

**Authors:** Xuejing Wang, Pinjia Que, Gerald Heckel, Junhua Hu, Xuecong Zhang, Chung-Yu Chiang, Qin Huang, Simin Liu, Jonathan Martinez, Nan Zhang, Emilio Pagani-Núñez, Caroline Dingle, Leung Yu Yan, Tamás Székely, Zhengwang Zhang, Yang Liu

## Abstract

Speciation with gene flow is an alternative to the nascence of new taxa in strict allopatric separation. Indeed, many taxa have parapatric distributions at present. It is often unclear if these are secondary contacts, e.g. caused by past glaciation cycles or the manifestation of speciation with gene flow, which hampers our understanding of how different forces drive diversification. Here we studied genetic, phenotypic and ecological aspects of divergence in a pair of incipient species, the Kentish (*Charadrius alexandrinus*) and the white-faced Plovers (*C. dealbatus*), shorebirds with parapatric breeding ranges along the Chinese coast. We assessed divergence based on molecular markers with different modes of inheritance and quantified phenotypic and ecological divergence in aspects of morphometric, dietary and climatic niches. These analyses revealed small to moderate levels of genetic and phenotypic distinctiveness with symmetric gene flow across the contact area at the Chinese coast. The two species diverged approximately half a million years ago in dynamical isolation and secondary contact due to cycling sea level changes between the Eastern and Southern China Sea in the mid-late Pleistocene. We found evidence of character displacement and ecological niche differentiation between the two species, invoking the role of selection in facilitating divergence despite gene flow. These findings imply that the ecology can indeed counter gene flow through divergent selection and thus contribute to incipient speciation in these plovers. Furthermore, our study highlights the importance of using integrative datasets to reveal the evolutionary history and underlying mechanisms of speciation.

## 1 INTRODUCTION

Understanding how strongly the evolutionary processes, i.e. selection, gene flow and genetic drift shape the divergence of closely related species has been a long-standing interest in evolutionary biology (Coyne & Orr 2004; Feder et al. 2013; Nosil & Feder 2012; Wagner et al. 2012; Wolf & Ellegren 2017). Allopatric speciation is conventionally considered as the prevalent mode of speciation in which physical barriers completely restrict gene flow between two populations, facilitating the initiation of divergence through genetic drift or selection (Bernardi et al. 2008; Carson & Clague 1995; Mayr 1963). If populations remain isolated for a period long enough after divergence has been established, then this divergence could be maintained even in the presence of gene flow after secondary contact (Le Moan et al. 2016; Zhou et al. 2016).

An increasing number of studies have shown that divergence can arise and be maintained due to selection imposed by heterogeneous environments despite the constraining effect of gene flow (Martin et al. 2013; Morales et al. 2017). Under this scenario, divergent selection or ecologically-mediated sexual selection operating on certain (“magic”) traits can lead to reproductive isolation, and incompatibilities in a few “speciation genes” may be enough to constrain the homogenizing effect caused by gene flow even at an early stage of speciation (Fitzpatrick et al. 2015; Nosil & Schluter 2011; Slatkin 1973). For instance, different host plant preferences result in different digestive and physiological traits in *Timema* walking-stick insects (Nosil 2007). Locally-adapted phenotypes in body size are related to swimming and foraging ecology in guppies (*Poecilia reticulata*), which can maintain population diversification in the face of extensive gene flow (Fitzpatrick et al. 2015). Though evidence of speciation with gene flow is accumulating (Martin et al. 2013; Morales et al. 2017; Schluter 2009; Seehausen et al. 2014), to what extent a balance between selection and gene flow can drive divergence remains largely unknown. This might be interesting in populations with high propensity of range-wide gene flow such as in waterfowl (e.g. Liu et al. 2012) and shorebirds (Eberhart□Phillips et al. 2015; Küpper et al. 2012).

Shorebirds are a group of migratory species with remarkable movement ability. Seasonal migration may increase the probability of individual dispersal between populations (Arguedas & Parker 2000; Procházka et al. 2011) and consequently drive frequent gene flow between geographically distant populations (Peters et al. 2012; Winker et al. 2013). Direct evidence using tracking approaches confirmed that extremely long-distance gene flow can be mediated through individual movements among remote breeding colonies of pectoral sandpipers (*Calidris melanotos*) (Kempenaers & Valcu 2017). Range-wide phylogeographic studies also revealed extensive gene flow in several migratory shorebird species (e.g. Trimbos et al. 2011; Küpper et al. 2012; Miller et al. 2014). This may result in weak genetic structure across a species and consequently prevent population-level divergence (Eberhart□Phillips et al. 2015; Jackson et al. 2017).

In this study, we test the role of ecology and gene flow on the divergence of two shorebird species, the Kentish Plover (*Charadrius alexandrinus*) and the white-faced Plover (*C. dealbatus*) (Figure 1). *C. alexandrinus* breeds in coastal areas and inland lakes in Europe, Asia and North Africa (del Hoyo et al. 2016). A previous study uncovered low genetic differentiation of C. alexandrinus across the Eurasian continent, and also between continental and island populations in East Asia (Küpper et al. 2012). C. dealbatus was formerly regarded as a subspecies of Kentish plover (Hartert & Jackson 1915; Swinhoe 1870). Breeding sites of *C. dealbatus* were documented along the coast of China from Fujian to Hainan Island, and in south middle Vietnam (BirdLife International 2016), yet its geographic range is uncertain (Kennerley *et al.* 2008). Previous studies have found subtle but diagnosable differences in morphometric, plumage and behavioral traits between the two taxa (Figure 1d), leading to the erection of *C. dealbatus* as a full species (Bakewell & Kennerley 2008; Kennerley et al. 2008; Rheindt et al. 2011). The first genetic investigation on the relationship between these two taxa provided no evidence of genetic differentiation and concluded that these species may only differ in a few genomic regions (Rheindt et al. 2011). Although based on a handful of genetic loci, this finding is noteworthy as it raises a probability of divergence in the face of gene flow between these two young species.

**FIGURE 1.**
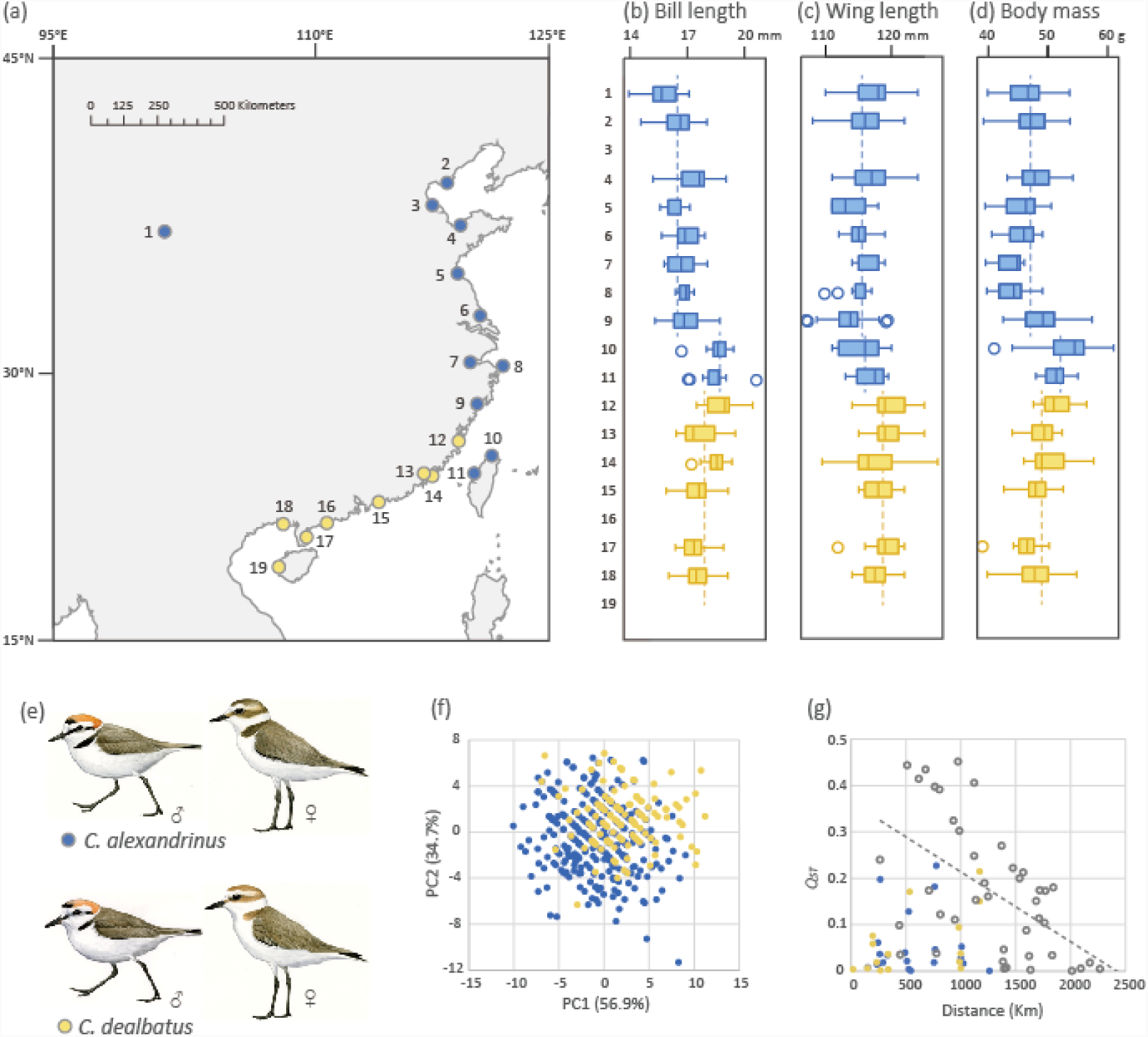
Sampling localities, morphology and trophic level differentiation of the Kentish Plover *Charadrius alexandrinus* (blue) and the White-faced Plover *C. dealbatus* (yellow). **(a)** Samples were from an inland site (1, Qinghai Lake), 16 Chinese mainland coastal localities (2, Tangshan; 3, Cangzhou; 4, Weifang; 5, Lianyungang; 6, Nantong; 7, Ningbo; 8, Zhoushan; 9, Wenzhou; 12, Fuzhou; 13, Xiamen; 14, Jinmen; 15, Shanwei; 16, Yangjiang; 17, Zhanjiang; 18, Beihai and 19, Dongfang), and two localities on Taiwan Island (10, Xinbei and 11, Zhanghua). **(b-f)** Differences between two species in several characters. *C. alexandrinus* has on average shorter bill length (except for Taiwan populations) **(b)**, wing length **(c)** and lower body mass (except for Taiwan populations) **(d)** than *C. dealbatus*. Localities with less than five measured adults were excluded in these analyses. **(f)** Plots of the first two principal components and their associated variance explained which showed the subtle morphometric differences between the two species. **(e)** *C. alexandrinus* has overall more darkish plumage than *C. dealbatus* and males of the latter species have less black tinged on face and neck during breeding season. **(g)** Pairwise morphological difference *Q*_*ST*_ plotted against geographical distance between coastal populations of *C. alexandrinus* and *C. dealbatus*. *Q*_*ST*_ values within species were marked by blue or yellow dots. *Q*_*ST*_ between species (grey circles) declined as the distance increased (regression showed as the dashed line).

Here we provide comprehensive data from multiple sources to explore divergence patterns between these two species of plovers. We carried out extensive sampling on breeding populations covering the potential area of contact along the Chinese coast. We obtained genetic polymorphism data from multiple markers with various mutation rates, i.e. mitochondrial DNA, exons and autosomal microsatellites. Using these new datasets, it is possible to estimate the intensity and direction of gene flow (Hey & Nielsen 2004; Won & Hey 2005) as well as other important demographic parameters such as the effective population size (*N*e) and the timing of divergence (reviewed in Hey 2006) between *C. alexandrinus* and *C. dealbatus*. Because it is difficult to directly quantify divergent selection, inference from the comparisons of traits which are potentially targets of selection are necessary to offer hints on selective mechanisms in maintaining divergence (Hoskin & Higgie 2010; Lande 1982; Nosil 2007). By collecting data on morphology, diet, and environmental niche, we attempt to 1) characterize geographic variation in genetic and phenotypic traits in the two plovers, 2) estimate demographic history and intensity of gene flow, and 3) infer the role of multiple factors, i.e. geographical premating isolation and gene flow during divergence between these two taxa. Taken together, our investigation provides new insights into the evolutionary history of two incipient bird species.

## 2 MATERIALS AND METHODS

### DNA sample collection

We mainly collected samples along the eastern coastal area of China (Fig 1a), and also obtained samples from the two biggest continental islands, Taiwan and Hainan, as well as two small islands that are close to the Chinese coastal line (Zhoushan and Jinmen). Sampling sites were separated by approximately 200-250 km along a 2,300 km transect, spanning almost the entire Chinese coastline. Further, one high altitude population (breeding at 3 350 m a.s.l.) near Qinghai Lake was sampled as an inland outgroup. Breeding individuals were captured using a funnel-trap method described in Székely et al. (2008) between March and July in 2014-2015. For each nest, blood samples of the breeding pair or a chick were collected. Tissue samples were also obtained from dead individuals found in the field. Species identification during sampling was based on the summary of plumage characters, morphometric, and ecological differences between C. *alexandrinus* and C. *dealbatus*, as described in Kennerley et al. (2008). Overall, we collected samples from 454 individuals from 19 breeding sites. All bird-capture and sampling were performed with permission from the respective authorities (Mainland China: Beijing Normal University to PJQ and Sun Yat-sen University to YL; Taiwan: Changhua County Government, New Taipei City Government and Kinmen County Government to ZYJ).

### Morphometric analysis

For each adult individual, four basic morphometric measurements were taken: body mass, wing length, tarsus length and bill length. Body mass was measured with an electronic scale (± 0.01 g). Wing length (flattened) was measured with a wing ruler (± 1 mm). Bill length to skull and tarsus lengths were measured using vernier callipers (± 0.01 mm). Measurements were taken by QH, PJQ and ZYJ following the standard described in Redfern & Clark (2001). To avoid potential biases, some individuals were measured twice or three times among these authors to make sure there was no significant difference (*p* < 0.05, between 20 – 30 trails) among them at the beginning and end of the fieldwork in the year 2014-2015.

We carried out principle component analysis (PCA) for the four traits to visualize the variation of morphometric data. Analyses of covariance (ANCOVAs) were conducted based on the PCA results to test statistical significance. We carried out t-tests between species for each measurement. Pairwise morphological difference *Q*_*ST*_ based on PC1 scores from PCA were calculated following Storz (2002) and plotted against linear distances between coastal populations. All aforementioned analyses were performed in PAST 3.12 (Hammer et al. 2001).

### DNA Sequencing and microsatellite genotyping

We extracted genomic DNA using Tiangen Blood & Tissue Genome DNA Kits, following manufacturer protocols (Tiangen, Beijing, China). DNA quality was measured with NanoDrop 2000 (Thermo Scientific, USA). Three mtDNA loci, partial ATPase subunit 6/8, partial D-Loop of the mitochondrial control region (CR) and NADH dehydrogenase subunit 3 fragment (ND3), were amplified for all samples using primers from Rheindt *et al.* (2011). PCR reactions for mtDNA amplification were carried out in 20 μl volumes containing 1X PCR buffer (Takara Shuzo, Japan), 10-50ng DNA template, 0.5 unit Taq DNA Polymerase (Takara Shuzo, Japan), 1.0 μM of each primer, 2.0 mM of each dNTP and 1.0 μM MgCl_2_. We also sequenced 14 autosomal and two Z-linked exonic loci (designed from Liu *et al*. 2018) for a subset of 40 individuals representing both species and several populations within species. PCR for nuclear loci had higher concentration of dNTP at 4.0 mM and 2.5 μM for MgCl_2_ and PCR cycling profiles are listed in Table S3. Each PCR product was checked on a 1% agarose gel and then sequenced on ABI3730XL (Applied Biosystems, USA) served by MajorBio, Shanghai, China.

We genotyped all individuals at 22 autosomal microsatellite markers (mst) mostly from Küpper *et al.* (2007) but with C204 from Funk *et al.* (2007) and Hru2 from Primmer *et al.* (1995). Microsatellite loci were amplified using three multiplex PCRs with respective cycling profiles (Table S3). Each PCR was in a total volume of 15 μl containing 1X HotStart buffer (Tiangen), 10ng DNA template, 1 unit Multi HotStart DNA Polymerase (Tiangen), 0.4 μM of each fluorescently labeled primer, 2.0 mM of each dNTP and 2.0 μM MgCl_2_. Multiplex PCR products and associated genotypes were isolated on ABI3730XL (Applied Biosystems, USA) served by Invitrogen, Shanghai, China and their length was determined using GeneMapper software v.3.7 (Applied Biosystems) against an internal size standard (GeneScan-500LIZ; Applied Biosystems).

### Genetic diversity and population structure analyses

For DNA sequence data, we aligned each mitochondrial or nuclear locus using the CLUSTAL W algorithm implemented in MEGA v.6.06 (Tamura et al. 2013), and the alignment was checked by eye and manually edited if needed. For nuclear DNA sequence data, we first used PHASE 2.1.1 (Stephens et al. 2001) to reconstruct haplotypes of nuclear sequences with heterozygous sites. Each run was set to 10,000 iterations, 100 burn-in and 10 thinning intervals. For both mtDNA and nuclear loci, basic genetic polymorphism statistics such as haplotype number *h*, haplotype diversity *Hd*, number of segregating sites *S*, nucleotide diversity π and Tajima’s *D* (Tajima 1989) of each locus and each population were calculated in DnaSP 5.10.1 (Librado & Rozas 2009). Haplotype networks of each locus were constructed using a median-joining algorithm (Bandelt et al. 1999) in PopART 1.7.2 (Leigh & Bryant 2015).

We used FreeNA (Chapuis & Estoup 2007) to check for the frequency of null alleles at each microsatellite locus. Further tests for Hardy-Weinberg equilibrium and pairwise linkage disequilibrium (LD) were carried out with Arlequin 3.1.1 (Excoffier et al. 2005). Hardy-Weinberg equilibrium tests were run with 10,000 permutations. LD tests were run for 100,000 steps of Markov chain. To obtain genetic diversity estimates, we calculated observed heterozygosity (*H*o) and expected heterozygosity (*H*_E_) in GenAlEx 6.5.1 (Peakall & Smouse 2012).

We estimated population structure between species and among sampling sites within species using several approaches. First, for mtDNA and microsatellites, we performed analyses of molecular variance implemented (AMOVAs) in Arlequin to assess the proportion of genetic variance explained by the different partition settings, e.g. (1) the two species, (2) *C. alexandrinus* continental populations and Taiwan island populations, *C. dealbatus*, (3) coastal populations (including Hainan Island), Qinghai and Taiwan. Second, we also calculated pairwise *Φ*_ST_ and *F*_ST_ between breeding sites for mtDNA and mst, respectively, and we derived significance levels using 10,000 permutations in Arlequin.

For microsatellite genotypes only, we carried out assignment analyses with two model-based Bayesian approaches. First, we performed a Bayesian clustering analysis using the admixture model with correlated allele frequencies implemented in STRUCTURE 2.3.4 (Hubisz et al. 2009; Pritchard et al. 2000). Ten independent analyses were run from *K*=1 to *K*=8 for 500,000 Markov chain Monte Carlo (MCMC) generations with 100,000 burn-in. Replicate runs were combined using STRUCTURE Harvester 0.6.94 (Earl 2012) and CLUMPP 1.1.2 (Jakobsson & Rosenberg 2007). The most likely number of genetic clusters was also determined using Structure Harvester using the criteria described in Evanno et al. (2005). Second, we used a Bayesian clustering algorithm that takes the geographical coordinates of each sampling site into account using the R package GENELAND 4.0 (Guillot et al. 2005). 1,000,000 MCMC iterations were run with thinning set to 100 from 1 to 10 populations, with maximum rate of Poisson process set to 100, uncertainty of spatial coordinates set to 0, maximum number of nuclei in the Poisson–Voronoi tessellation to 300, and independent Dirichlet distribution model for allele frequencies. With the same package, the most likely number of clusters was determined based on their posterior density. To confirm the consistency of the results, we repeated the MCMC simulation 10 times.

Furthermore, to determine the probability that an individual was a hybrid or a migrant, we estimated the posterior probability of each individual based on multilocus genotypes. First we used HYBRIDLAB 1.0 (Nielsen et al. 2006) to simulate 100 individuals of each species, F1, F2 and back-crosses in each direction from microsatellite genotypes of 260 individuals with q-value higher than 95% in STRUCTURE analysis. 100 simulated parents from each species and 100 F1 were used to calculate the threshold for individual hybrid assignment in STRUCTURE following Vähä and Primmer (2006). Finally, NewHybrid 1.1 (Anderson & Thompson 2002) was used to identify potential hybrid individuals. 100,000 burn-in followed by 400,000 sweeps were performed for both simulated and real data.

### Demographic analysis

To infer the demographic histories of the two plover species, we applied Isolation with Migration model (IM) (Hey & Nielsen 2007) analyses using the combined sequence data set of 16 nuclear and three mtDNA loci based on 20 individuals of each species. IM analysis allows the inference of genealogies under different demographic scenarios and estimation of population genetic parameters, such as divergence times, effective population sizes and migration rates between species since their divergence from the common ancestor. The homologues Killdeer *Charadrius vociferus* sequences from GenBank were used as outgroup. The substitution rate of each locus was calculated using the method in Li et al. (2010). The ratio of net genetic distance of each locus across ingroup–outgroup was calculated, compared with net distance of mitochondrial cytochrome b (*cytb*) and then multiplied by the substitution rate for *cytb* (0.0105 ± 0.0005 substitution/site/mya, Weir & Schluter 2008). We used the Phi test (Bruen et al. 2006) for recombination within nuclear loci and no recombination was detected. We implemented models in IMa2p (Sethuraman & Hey 2016), the parallel version of IMa2 (Hey 2010). For each analysis, we ran 48 MCMC chains for 2,500,000 steps of burn-in followed by 500,000 genealogies saved, each recorded after 100 steps. Because IM analysis only provides estimates of average gene flow since the divergence of the two species, we used a Bayesian analysis in BayesAss 3.0.4 (Stephens & Donnelly 2003) based on microsatellite genotypes to characterize the level of gene flow within recent generations. We performed 10,000,000 iterations and 1,000,000 burn-in.

### Ecological niche modeling and niche overlaps

To infer potential past range shifts induced by climatic changes, we carried out ecological niche modeling (ENM) using Maxent 3.3.3k (Phillips et al. 2009). The occurrence records of the two species of plovers were obtained from online databases of the Global Biodiversity Information Facility (GBIF, http://www.gbif.org/), China Bird Report (http://www.birdreport.cn) and our records during the sampling expeditions. Further, eight bioclimatic variables (i.e., the mean diurnal range, isothermality, minimum temperature of the coldest month, mean temperature of the warmest quarter, annual precipitation, precipitation of the driest month and precipitation seasonality) were obtained from the WorldClim database v.1.4 (Hijmans et al. 2005).

To explore niche similarity between the two species, we performed an ordination null test of PCA-env in environmental (E)-space (Broennimann et al. 2012; Hu et al. 2016). The PCA-env calculates the occurrence density and environmental factor density along environmental (principal component) axes for each cell using a kernel smoothing method and then uses the density of both occurrences and environmental variables to measure niche overlap along these axes (Broennimann et al. 2012). An unbiased estimate of Schoener’s D metric was calculated for our data using smoothed densities from a kernel density function to measure niche overlap between the two species that is ensured to be independent of the resolution of the grid. Statistical confidence in niche overlap values was then tested through a one-sided niche-similarity test (Broennimann et al. 2012). All statistical analyses were performed in R 3.0.2 (R Development Core Team 2013) using scripts available in Broennimann et al. (2012). Details of ENM construction and niche-similarity tests are available in Appendix S1.

### Dietary niche differentiation inferred by stable-isotope analysis

Because stable isotopic compositions of consumer tissues can be used to estimate the relative contribution of assimilated dietary sources (DeNiro & Epstein 1978), stable-carbon(C) and nitrogen(N) isotope analysis is widely used as a tool to study avian dietary patterns (Hobson & Clark 1992; Pagani-Núñez et al. 2017). Carbon isotope ratios differ between C3, C4 and CAM plants due to differences in the photosynthetic pathways, and these differences are incorporated into an animal when the plants are consumed and so can be used to infer information about dietary niches (Hobson & Clark 1992). N isotopes are useful for identifying species/individuals which occupy different trophic positions (high δ^15^N implies higher trophic level; Chen et al. 2017). In order to compare dietary differences based on differences in δ^15^N and δ^13^C between the two species, we collected the outer pair of rectrices from seven adults per site at eight sites: Qinghai Lake, Tangshan, Lianyungang and Rudong for *C. alexandrinus*; Fuzhou, Shanwei, Zhanjiang, Dongfang for *C. dealbatus*. Since both species perform a complete post-breeding molt within their breeding grounds (Ginn & Melville 1983, personal observation), isotope ratios represent trophic level and habitat preferences during the breeding period. We estimated niche width and overlap per species using an isotopic Bayesian approach based on δ^13^C and δ^15^N profile. Detailed information on lab procedures and statistical analyses can be found in Appendix S2.

## 3 RESULTS

### Morphological differentiation

The two plover species showed subtle but significant differences for most morphological traits (ANCOVA; *p* < 0.001, Figure 1) even though there was considerable overlap in morphology between the species at the level of the individual (Figure 1f). On average, *C. dealbatus* had a longer bill (17.91 ± 0.94 mm vs. 16.69 ± 1.06 mm in *C. alexandrinus, p* < 0.001, Figure 1b), longer wings (118.58 ± 2.90 mm vs. 115.59 ± 3.02 mm in *C. alexandrinus, p* < 0.001, Figure 1c), and larger body mass (48.99 ± 3.26 g vs. 47.56 ± 3.73 g in *C. alexandrinus, p* = 0.001, Figure 1d, Table S5) than those of *C. alexandrinus*. There was no difference in tarsus length between the two species (*p* = 0.962). Individuals of *C. alexandrinus* from Taiwan Island were heavier than continental populations of *C. alexandrinus* and *C. dealbatus* (both *p <* 0.001). *Q*_*ST*_ between coastal populations of *C. alexandrinus* and *C. dealbatus* was negatively correlated with geographic distance (R^2^ =0.293, *p <* 0.001, Figure 1g).

### Genetic diversity and population structure

We sequenced 357 individuals at all three mtDNA loci (224 *C. alexandrinus* and 133 *C. dealbatus,* GenBank Accession No. xxxx-xxxx). For each site, sample size ranged from 11 to 30 individuals. For each individual, we obtained in total 1729 base pairs (bp) of mtDNA sequence, including 846 bp ATPase6/8, 505 bp D-loop and 378 bp ND3. *C. alexandrinus* showed higher intraspecific genetic diversity than *C. dealbatus* (Table 1). Haplotype networks show that in all loci, most individuals were sorted into two major haplogroups, corresponding to the two species of plovers. One non-synonymous substitution separated the two haplogroups at both ATPase6/8 and ND3 loci (Figure 2a). Moreover, a subset of samples containing 20 individuals of each species were sequenced at 16 loci (range 440-902 bp for each locus; Table 1 and Table S1, GenBank Accession No. xxxx-xxxx) for a total of 11,209 bp nuclear DNA sequence. The haplotype networks from autosomal and Z-linked loci did not show strong patterns of lineage sorting like mtDNA. The most common haplotypes were shared by both species of plover (Figure 2b and S1). Moreover, both species showed the signal of recent demographic expansion as detected by significant Tajima’s *D* values (Table 1).

**TABLE 1.**
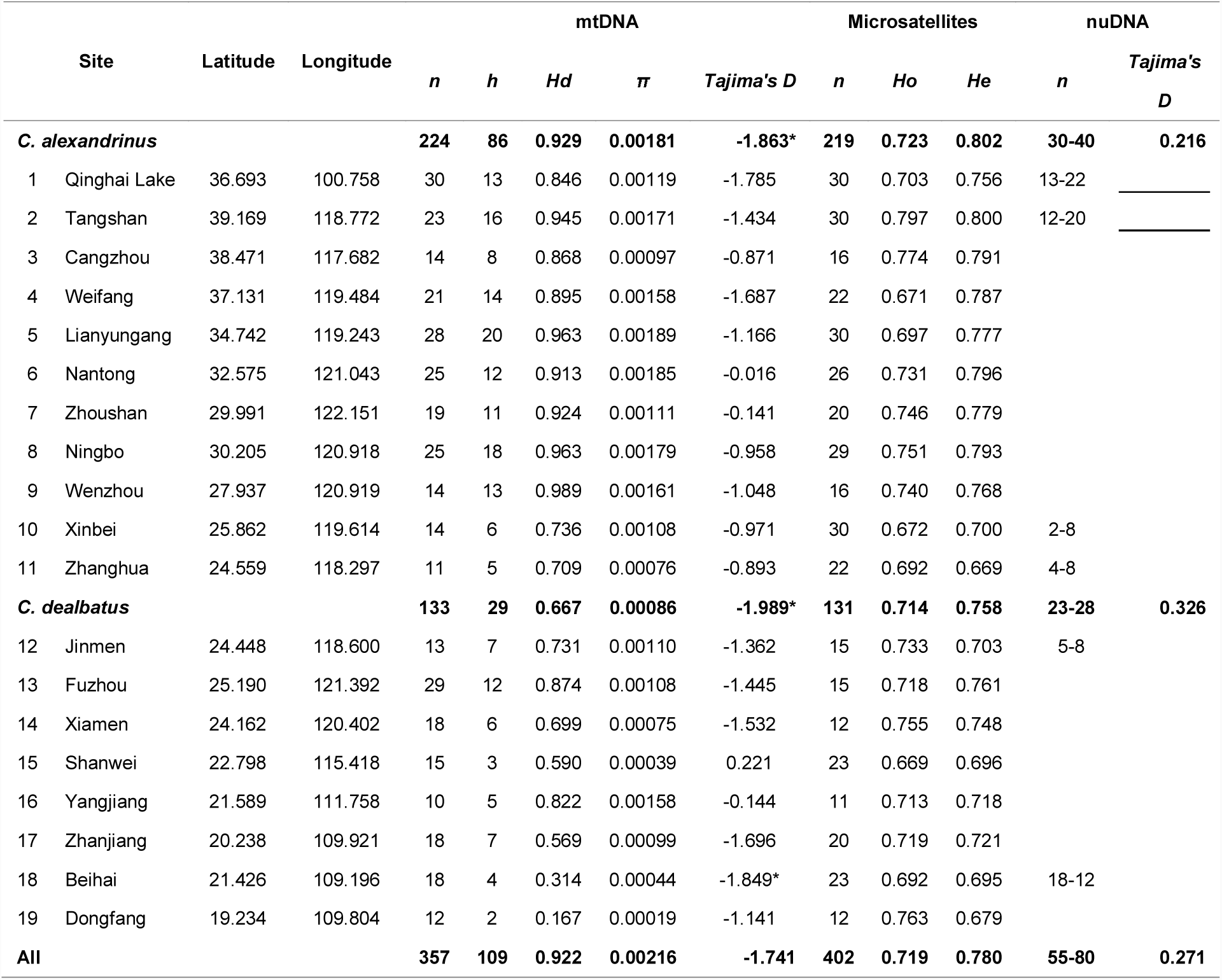
Sampling localities and genetic polymorphism of *C. alexandrinus* and *C. dealbatus*. The number of individuals (*n*) analyzed for mtDNA, autosomal microsatellites and nuclear exonic loci (nuDNA) are given. Site number corresponds to the numbers in Figure 1. Estimates of *h*, number of haplotypes; *Hd*, haplotype diversity; π, nucleotide diversity; *Tajima’s D* value, *Ho*, observed heterozygosity; *He*, expected heterozygosity were calculated for each locality and species.

**FIGURE 2.**
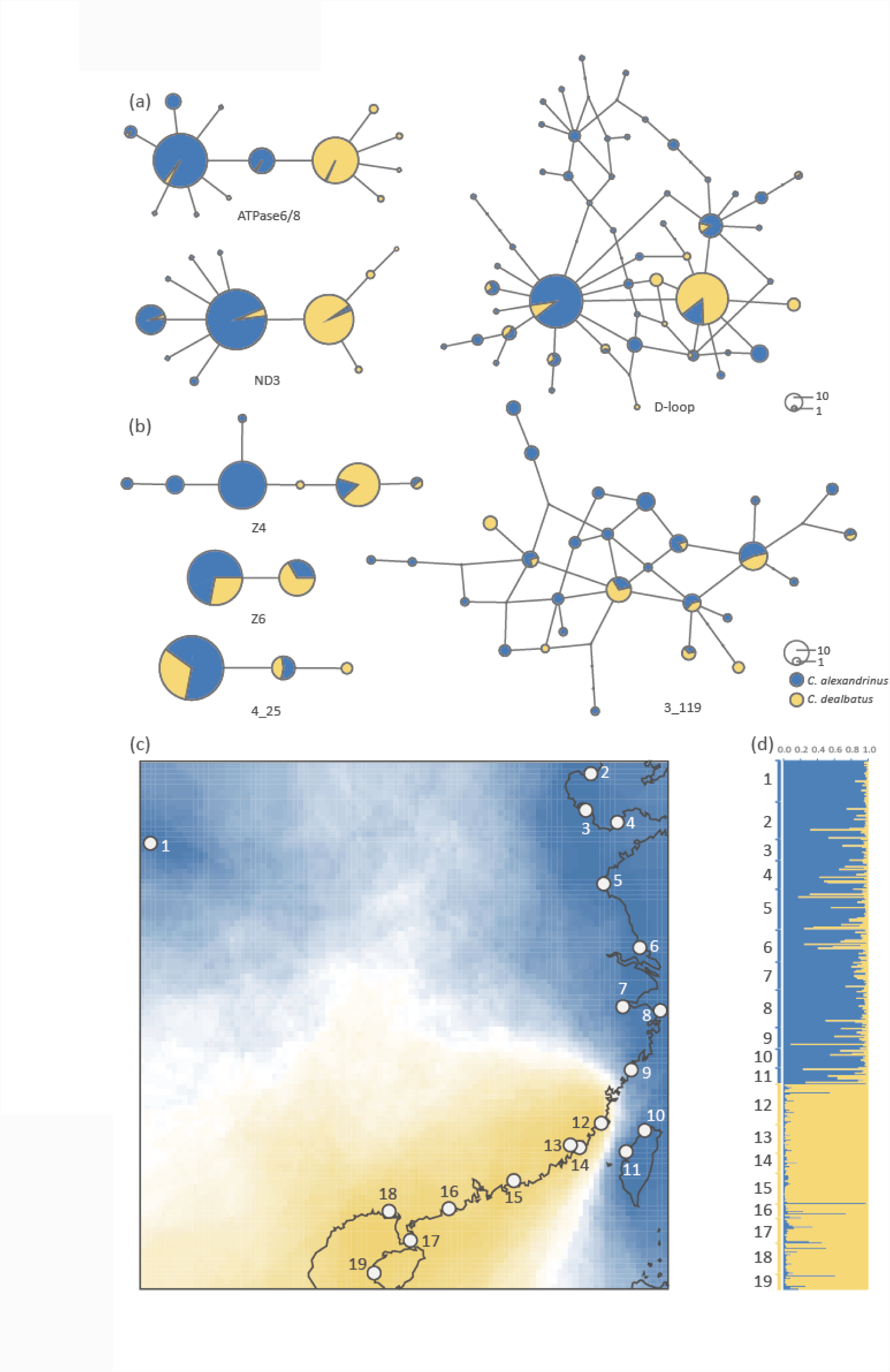
Haplotype networks based on mitochondrial and nuclear DNA sequences, and population genetic structures based on microsatellite loci. *C. alexandrinus* marked in blue and *C. dealbatus* in yellow. **(a)** Haplotype networks of three mitochondrial loci. For ATPase6/8 and ND3, over 95% of the individuals are sorted into two major haplogroups. **(b)** Examples of haplotype networks based on 55 to 80 individuals of four exonic nuclear loci. **(c)** Genetic clustering inferred with Geneland based on microsatellite genotypes. Blue and yellow show the assignment probability of a location for alternative genetic clusters. Location numbers are consistent with Figure 1. **(d)** Genetic clustering inferred with STRUCTURE using microsatellites.

Overall genetic differentiation was significant and high in mtDNA data (*Φ*_ST_ = 0.506, *p* < 0.001) and low at microsatellite loci (*F*_ST_ = 0.036, *p* < 0.001). For nuclear sequence data, genetic differentiation between species was also significant at autosomal loci (*Φ*_ST_ = 0.100, *p* < 0.001) and particularly high at the Z-linked loci Z4 (*Φ* _ST_ = 0.726, *p* < 0.001) and Z6 (*Φ*_ST_ =0.309, *p* < 0.001, Table S3). In the AMOVA analysis, we observed the largest difference between groups when the data were partitioned by species (i.e. *C. alexandrinus* and *C. dealbatus*), with this grouping explaining 49.7% of the variance in mtDNA and 2.4% in the microsatellites (*p* < 0.001; Table 2). These were significantly higher than the values of genetic variation when we partitioned samples as coastal vs. island populations or coastal population vs. Qinghai vs. Taiwan populations. Within *C. alexandrinus*, minor genetic differentiation was found between the inland population (Qinghai Lake) and coastal populations (mtDNA *Φ*_SC_ = 0.042, *p* = 0.022; mst *F*_SC_ = 0.021, *p* < 0.001). *C. alexandrinus* populations on Taiwan Island also shared haplotypes with other coastal populations (mtDNA *Φ*_SC_ = 0.021, *p* =0.146); but were significantly differentiated in microsatellites (microsatellite *F*_SC_ = 0.016, *p* < 0.001).

**TABLE 2.** Hierarchical analyses of molecular variance (AMOVA) based on concatenated mtDNA data and 13 microsatellite loci for *C. alexandrinus* and *C. dealbatus*. Samples were partitioned in three groupings: 2 groups (*C. alexandrinus* and *C. dealbatus*); 3 groups (*C. alexandrinus* continental populations and Taiwan island populations, *C. dealbatus*); 4 groups (*C. alexandrinus* inland (Qinghai Lake), *C. alexandrinus* coastal, *C. alexandrinus* Taiwan island, and *C. dealbatus*). *Va*, genetic variation among groups; *V*b, variation among populations within groups; *V*c, variation within populations. Between-group genetic differentiation was highest when populations were partitioned into two groups (*C. alexandrinus* and *C. dealbatus*).

|  | Grouping | *Va*% | *Vb*% | *Vc*% |
| --- | --- | --- | --- | --- |
| **mtDNA** | **2** | **49.712** | **8.491** | **41.797** |
|  | 3 | 45.695 | 9.269 | 45.036 |
|  | 4 | 41.782 | 10.296 | 47.922 |
| **microsatellites** | **2** | **2.416** | **1.798** | **95.786** |
|  | 3 | 2.192 | 1.749 | 96.059 |
|  | 4 | 1.917 | 1.776 | 96.307 |

All $p < 0.001$

For microsatellite loci, 18 out of 22 markers were successfully genotyped. However, four markers (Calex-04, 08, 19, C204) showed a large proportion of missing data, while another four loci (Calex-11, 24, 26, 43) showed an estimated frequency of null alleles over 10% and moreover one locus (Calex-35) showed H-W disequilibrium (Table S2). Consequently, the aforementioned nine loci were removed from the dataset, making genotypes at 13 loci of 402 (271 *C. alexandrinus* and 131 *C. dealbatus*) individuals for the final microsatellite dataset for further analysis.

The results of the STRUCTURE analysis clearly showed two genetic clusters representing each species (Figure 2d) and no obvious gradual transition along the coastline that could be expected from a hybrid zone. The average *Delta K* value when *K* = 2 was much higher than other options (Figure S2). Using georeferenced data in GENELAND, our results corroborated the genetic clustering patterns inferred from STRUCTURE. The posterior probability was 0.70 when *K* = 2 in contrast with the posterior probability of 0.25 when *K* = 3. We visualized the GENELAND results in a map with the probability distribution of posterior mode of class membership, which further supports the separation between *C. alexandrinus* and *C. dealbatus* along the China coast (Figure 2c). The divide between these two species was located between Wenzhou and Fuzhou according to the GENELAND results, but it is unclear if these two species come into contact or forms a putative hybrid zone in this region. Again, the individuals from the sites in Taiwan were assigned to the cluster of *C. alexandrinus* (Figure 2c).

### Demographic history in two plovers

Isolation with migration analyses suggested that *C. alexandrinus* and *C. dealbatus* diverged 0.56 (0.41 – 5.19) million years ago. Estimated migration rates in both directions were significant (*p* < 0.001) with slightly higher gene flow from *C. alexandrinus* into *C. dealbatus* (2.69 migrants per generation) than the vice versa (ca. two migrants per generation). The estimated effective population size of *C. alexandrinus* (N_e_ ≈ 1.59 – 4.44 million) was about 8-10 times higher than for *C. dealbatus* (N_e_ ≈ 0.15 – 0.52 million).

The estimation of recent gene flow between the two species with BayesAss using microsatellites also suggested bidirectional gene flow. Gene flow from *C. dealbatus to C. alexandrinus* was slightly higher (0.013, *p* = 0.028) than that in the other direction (0.010, *p* = 0.028). In *C. alexandrinus* populations, one likely migrant from *C. dealbatus* and two hybrid F1 individuals were identified (probability higher than 0.5). With the same threshold value, two migrants and one F1 individual were identified in *C. dealbatus* populations.

### The detection of hybrid individuals

Analyses with NewHybrids provided no evidence of a large number of hybrids or a higher frequency of these near the potential contact area between the species. Based on the STRUCTURE results and simulations, the optimal threshold for distinguishing hybrids from non-hybrids was q=0.836. Based on this threshold, 81.9% of the individuals (204 out of 246) collected from sites at the northern Chinese coastline, Qinghai Lake and Taiwan Island were assigned to *C. alexandrinus* (Figure 2c-d). Individuals with intermediate q-values, which were possibly hybrids, were found at most northern Chinese sampling sites (Figure 2d). For the southern coastline in China, 145 out of 156 Individuals (92.9%) belonged to *C. dealbatus* with a probability larger than 0.836 (Figure 2c-d). Only one individual of each species was assigned to the genetic cluster representing the other species with high probability (Figure 2d).

The result of NewHybrids was highly concordant with STRUCTURE results and suggested migrants moving in both directions. NewHybrids consistently identified three migrants mentioned above with probability higher than 99%. However, NewHybrids failed to identify simulated hybrid individuals and recognized them as migrants. Thus, the result of NewHybrids was not used for hybrid identification (Fig S3).

### Niche modeling, projections and comparisons

Our ecological niche models effectively captured the current distribution of both *C. alexandrinus* and *C. dealbatus* (Figure 4) with a high discrimination capacity (AUC values > 0.88 for training and test data). Jackknife tests on variable importance for *C. alexandrinus* revealed that isothermality, precipitation seasonality and mean temperature of the warmest quarter were the three highest ranked variables when used in isolation. For *C. dealbatus*, mean diurnal range and annual precipitation were the most important variables. The simulation of the three periods, i.e. Last Interglacial (LIG, 120-100Ka), Last Glacial Maximum (LGM, 21Ka, MIROC model) and current times, respectively, showed range shifts in both species. In particular, these results suggest that ranges of both species shrank during the LIG in the Chinese coastal area accompanied by an increase of climatic suitability in the inland region for *C. alexandrinus*. In contrast, suitable habitats expanded for both species during the LGM in the coastal area, the East China Sea and the northern part of the South China Sea (sea area between Hainan and Taiwan), probably due to the fall of the sea level.

**FIGURE 3.**
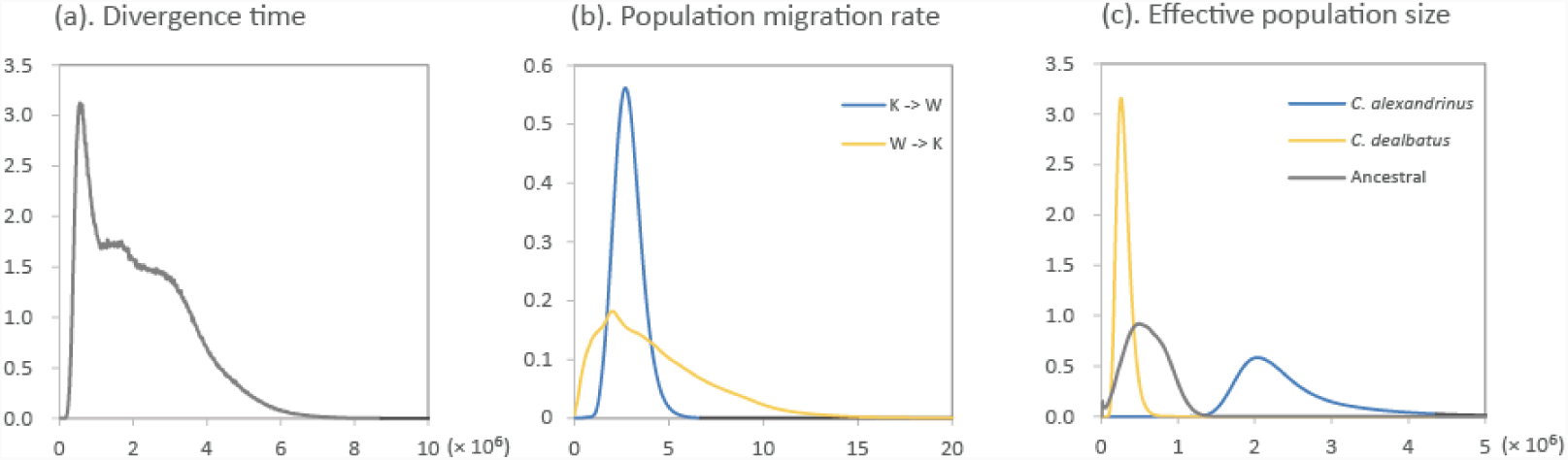
Posterior densities of population demographic parameters estimated using the isolation-with-migration model (IM) implemented in IMa2p. These analyses used the combined data set of three mitochondrial (1729 bp) and 15 nuclear exonic loci (11209 bp). K represents the Kentish Plover *C. alexandrinus*; W represents White-faced Plover *C. dealbatus*. **(a)** Population divergence times (*T*) of *C. alexandrinus* and *C. dealbatus*. **(b)** Population migration rates (*2NM*) stand for the average number of individuals in each group that were migrants from the other group in the past. Gene flow on both directions was significant. **(c)** Effective population sizes (*Ne*) of the two species and their most recent common ancestor.

**FIGURE 4.**
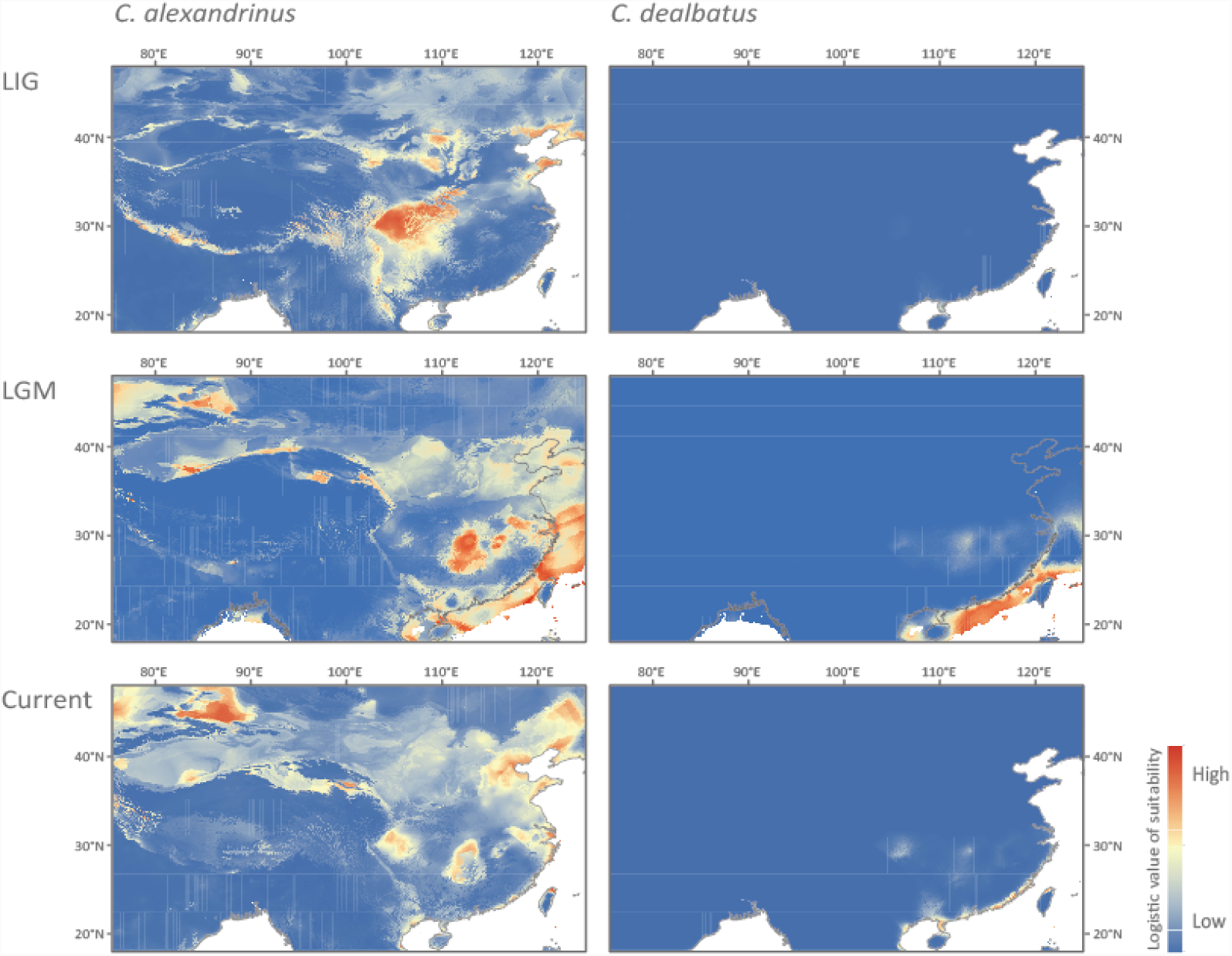
Predicted environmental suitability for *C. alexandrinus* (left) and *C. dealbatus* (right), via ecological niche modeling (ENM). ENM results are shown for the Last Interglacial (LIG, 120-100Ka), Last Glacial Maximum (LGM, 21Ka, MIROC model) and current times, respectively.

The ordination approach using PCA-env suggested that the overlap of the current climatic niches of the two species is relatively low (Figure S4a-b). The two species can be separated based on the first two PCs with an accumulative 81.5% of the total variance explained (Figure S4c). The niche equivalency test rejected the null hypothesis that the species pair is distributed in identical environmental space (*p* = 0.019; Figure S4d).

### Stable-isotope analysis

Our stable-isotope analysis showed that *C. dealbatus* exhibited significantly higher δ^15^N values than *C. alexandrinus* (*p* < 0.001, Figure 5a, Figure S5). In contrast, we did not find significant differences in δ^13^C between the two species (*p* = 0.161). Further isotope space overlap analysis showed that *C*. *alexandrinus* individuals had a high probability to be found within the isotopic niche space of *C. dealbatus* (95.3%), while *C. dealbatus* individuals showed a relatively low probability to be found in the isotopic niche space of *C*. *alexandrinus* (46.5%, Figure 5b). Moreover, we found that *C. alexandrinus* showed higher variability in δ^15^N profile across breeding populations than *C. dealbatus.*

**FIGURE 5.**
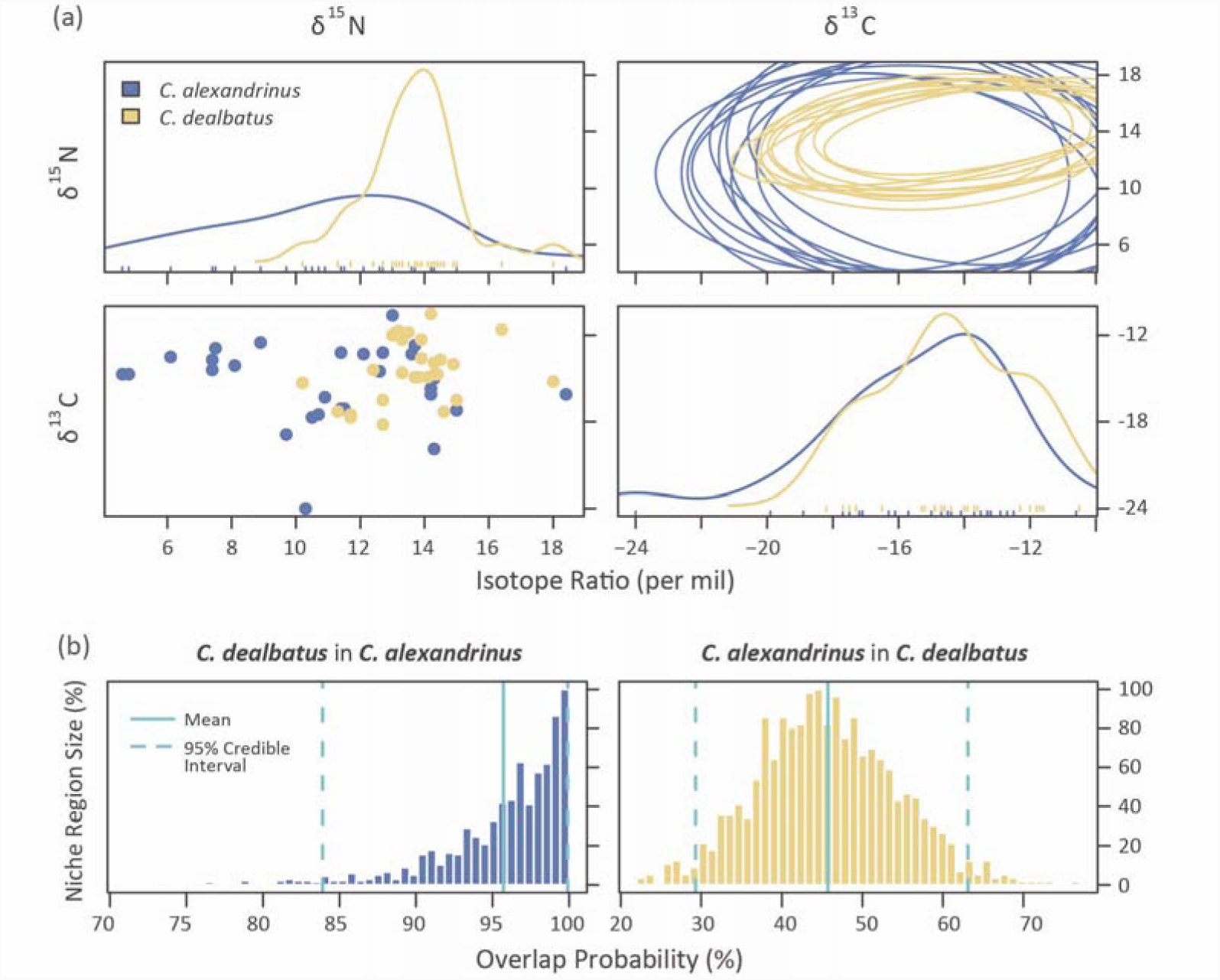
**(a)** Raw data (bottom-left), density distribution (top-left and bottom-right), and stable isotope related to dietary niche regions (top right) of δ^13^C and δ^15^N of *C. alexandrinus* and *C. dealbatus* feathers generated by “nicheROVER” (Swanson *et al*. 2015). The niche region of C. *alexandrinus* (blue) was broader than and contained that of *C. dealbatus* (yellow). **(b)** Distribution of the posterior probability that an individual from one species is found within the niche region of the other species. Vertical lines for mean and 95% credible intervals are included in the histogram of each overlap metric.

## 4 DISCUSSION

Our genetic data show that *C. alexandrinus* and *C. dealbatus* have diverged to a level of advanced sorting of alleles between the two species particularly in mtDNA and Z-linked genes with lower effective population sizes (Figure 2a-b). For autosomal microsatellite data, we also found that genetic differentiation is low at intraspecific level but substantially high between the species (Figure 2c-d, Table 2). Though it is unclear whether a narrow hybrid zone exists in the contact area on the Chinese coast, a considerable level of symmetric gene flow occurred between the two plovers (Table 3). We find diagnosable differences in morphometric traits and ecological characters between the two plovers along the coast. At odds with these results, a previous work found no evidence of genetic differentiation between the two plovers (Rheindt et al. 2011). The present dataset comprises of systematically sampling along the Chinese coast, a larger suite of genetic markers and well-characterized ecological traits (Figure 1) while Rheindt et al. (2011) analyzed samples collected outside the breeding season. Potentially erroneous species assignment of non-breeding birds may underestimate genetic differentiation, because the distinguishing features in non-breeding plumage are relatively subtle (Bakewell & Kennerley 2008; Kennerley *et al.* 2008; Rheindt et al. 2011) and non-breeding *C. alexandrinus* and *C. dealbatus* mix frequently on the coast of the South China sea (own observation and Kennerley et al. 2008).

**TABLE 3.** Posterior mode, mean and range of 95% highest probability distribution (HPD) of six demographic parameters inferred with IMa2p between *C. alexandrinus* and *C. dealbatus.* Divergence times (*T*) are given in million years ago (Ma). The effective population size (*Ne)* of *C. alexandrinus* was about eight times that of *C. dealbatus.* Migration rate (*2NM*) into each species was about the same. K represents Kentish Plover *C. alexandrinus*; W represents White-faced Plover *C. dealbatus*; A is the most recent common ancestor of two species.

| Parameter | *T*/Ma | *Ne<sub>W</sub>*/10<sup>6</sup> | *Ne<sub>K</sub>*/10<sup>6</sup> | *Ne<sub>A</sub>*/10<sup>6</sup> | *2NM<sub>K→W</sub>* | *2NM<sub>W→K</sub>* |
| --- | --- | --- | --- | --- | --- | --- |
| **Mode** | 0.557 | 0.259 | 2.036 | 0.495 | 2.696* | 2.005* |
| **Mean** | 2.160 | 0.297 | 2.450 | 0.589 | 2.859 | 4.306 |
| **2.5% HPD** | 0.413 | 0.154 | 1.590 | 0.134 | 1.581 | 0.528 |
| **97.5% HPD** | 5.187 | 0.515 | 4.442 | 1.098 | 4.503 | 11.500 |

\* *p* < 0.001

### Phylogeographic patterns in *C. alexandrinus* and *C. dealbatus*

Two genetic lineages were found among breeding *Charadrius* plovers along the Chinese coast, corresponding the northern lineage to *C. alexandrinus* and the southern one to *C. dealbatus*, respectively (Figure 1a, Table 2). The sharp genetic break between the two lineages lies between Wenzhou and Fuzhou (latitude 26-27 °N) north of the Taiwan Strait (Figure 2c-d). Furthermore, samples from Taiwan Island belong to *C. alexandrinus* but the population in Jinmen Island is affiliated with *C. dealbatus* (Figure 2c-d). This pattern likely reflects historical isolation of the two species through separation by some geographical barriers at the Chinese coast during the Pleistocene climatic fluctuation periods. Divergence between the two plovers 560,000 years ago was inferred based on IM for the two species (Table 3, Figure 3).

This period falls into the Marine Isotope Stage 16 of the mid-late Pleistocene, during which was one of the Ice Age (Railsback et al. 2015). Accordingly the divergence between the plovers was probably linked to vicariance when the coastline was separated by a land-bridge raised due to the shallow sills between the East and the South China Sea (Ni et al. 2014). *Charadrius* plovers have originated in the Northern hemisphere and then radiated to the Southern hemisphere as suggested by a recent molecular phylogeny (Dos Remedios et al. 2015). It is thereby likely that *C. dealbatus* originated from *C. alexandrinus* and could have been diverging along the east China coast since the mid-late Pleistocene.

Phylogeographic patterns have been relatively well characterized in shorebirds breeding at high latitudes (Trimbos et al. 2011; Küpper et al. 2012; Miller et al. 2014). Panmixia or weak genetic differentiation was usually suggested, most likely driven by extensive gene flow. However, it seems that islands can act as a barrier to natal dispersal for the continental Kentish plovers (Küpper et al. 2012). In contrast, population structures in temperate and subtropical shorebird populations are poorly documented, probably due to a low level of species diversity. To the best of our knowledge, our study shows the first documented phylogeographic break in a bird species in the coastline in China. Interestingly, concordant phylogeographic patterns have been described in several coastal marine taxa, such as plants (Wang et al. 2015), fishes (Ding et al. 2018; Liu et al. 2007), shellfishes (Ni et al. 2012), and crustaceans (Wang et al. 2008). Though the exact splitting times are not congruent, the observed split line is at approximately 25°N latitude between the East China Sea and the Southern China Sea (reviewed in Ni et al. 2014). This consensus pattern resulted in the hypothesis that historical factors, i.e. sea level fluctuation during the Pleistocene, caused a convergent phylogeographic pattern in multiple coastal marine fauna in the marginal northwestern Pacific Ocean (Ni et al. 2014; Ding et al. 2018). Thus, our study contributes a vertebrate case to the accumulating literature about this species divergence hotspot. Apart from the major role of physical barriers, comparative phylogeographic studies also revealed that other abiotic factors, like ocean current and hydrothermal conditions, as well as species ecological characters, i.e. dispersal ability, habitat preference, life-history, and population demography can also play a role in contributing to divergence of coastline fauna (Liu et al. 2007; Ni et al. 2014).

### Morphological and ecological differentiation along a latitudinal gradient

The two plovers, *C. alexandrinus* and *C. dealbatus* are distributed along the Chinese coastline across a latitudinal and associated environmental gradients from temperate to tropical zones. Within each species, we found a general trend that northern populations have larger values than the southern counterparts in morphometry (Figure 1). This pattern may be related to Bergmann’s rule which states that body size of homeothermic animals is larger in colder climates than in warmer ones (Bergmann 1848). Populations of both plover species distributed in such a north-south gradient may benefit from an optimal temperature control (Salewski & Watt 2017). A mutually non-exclusive explanation is that the difference in morphological traits, i.e. body mass and bill length link to potential differences in resource exploitation and life history. Our data show that *C. dealbatus* has a larger average body mass than *C. alexandrinus.* Body mass is a comprehensive trait reflecting nutrition assimilation, energy reservation and expenditure (McNab 2009). The difference in body mass might be related to the difference in migration behavior where a lighter body mass in *C. alexandrinus* is favored by decreased transport cost of fuel storage during migration (Alerstam et al. 2003; McWilliams & Karasov 2001). In contrast, larger body mass in *C. dealbatus*, a short-distance migrant or resident, may be beneficial in multiple reproduction within a single breeding season while *C. alexandrinus* produces usually only a single clutch (Lin et al. in prep.). Difference in bill length may be driven by the difference in the use of food resources (Badyaev et al. 2008) but recently, the function of the bill as temperature regulator also started to attract scientific attention (Tattersall et al. 2016). On the other hand, an increasing number of studies have demonstrated that birds at higher latitude and in cooler environments have shorter bills (e.g. European sparrows, Johnston 1969; Australian shorebirds, Nebel et al. 2013), consistent with Allen’s Rule (Allen 1877). Nevertheless, we find that populations of *C. alexandrinus* in Taiwan had similarly large body mass and bill lengths as *C. dealbatus* populations (Figure 1b-d, g), which are significantly larger than mainland conspecific populations. This is probably caused by phenotypic plasticity in the Taiwan population to a sub-tropical ecology and resident life history, similar to the populations of *C. dealbatus* in the South China sea coast.

Climatic niche modelling indicates that the climatic conditions of the two species are spatially separated (Figure S4). *C. dealbatus* is restricted to breeding sites close to the coast, particularly in warmer tropical climate. In contrast, *C. alexandrinus* has a wider climatic niche, as represented by a broader climatic zone (Figure 4). This species can breed not only on temperate coasts (Que et al. 2015) but also in inland saline lake shore (Cramp & Perrins 1983), just as Qinghai Lake. In addition, our isotope analysis revealed that *C. alexandrinus* covered a wider range of isotope ratios than *C. dealbatus*, but *C. alexandrinus* exploited a lower trophic range (δ^15^N). This indicates that *C. dealbatus* probably feeds on a higher energy diet than its sister species (Hobson & Clark 1992). However, it is unclear whether such difference may result from diet preference divergence or due to differences in food resource availability (McCormack et al. 2010). Taken together, these results suggest that the ecological niches of the two plovers are significantly different in several aspects, and support a role for ecology in constraining range limits and perhaps habitat preference for the two shorebird species.

### Incipient speciation with ongoing gene flow

We find evidence of incipient speciation with ongoing gene flow between the two plovers. The IM results indicate a relatively young split with considerable gene flow between the two species (Figure 3 and Table 3). We suggest this is probably due to gene flow caused by secondary contact during the Pleistocene, because our niche modeling analysis reveals that the ranges of the two species were expanded and consequently overlapped between the Eastern and Southern China Sea during the Last Glacial Maximum (Figure 4). While our niche modeling demonstrates a cycle of range dynamics caused by sea-level changes, one should bear in mind that the last million years witnessed multiple glaciation cycles (Taylor et al. 2014, 2015). In particular, marine oxygen isotope records designate 28 isotope stages corresponding to glacio-climatic cycles in the last million years (Railsback et al. 2015). For ocean marginal and coastline taxa, this may imply that the fluctuations in sea level can lead to alternation between population isolation and contact throughout their evolutionary histories. This can in turn yield significant consequences on population divergence and historical demography because speciation can proceed through cycles of allopatric stages interspersed by parapatry, resulting in a diversification process characterized isolation with migration (Rheindt & Edwards, 2011).

Under this premise, a key question arises: after the two plovers diverged, how was their divergence maintained in the presence of ongoing gene flow? Speciation theory predicts that divergence is initiated either by genetic drift or divergent selection (Lynch et al. 2016; Rundle & Nosil 2005; Seehausen et al. 2014; Wolf & Ellegren 2017). However, genetic drift is unlikely to have been the only force to initiate divergence in the plovers for two reasons: the large effective population sizes (*Ne*) and a relatively high level of gene flow (Table 3). Moreover, large *Ne* and recent divergence make it unlikely that genetic drift could have led to diagnosable differences between populations (Ellstrand & Elam 1993; Lanfear et al. 2014), and gene flow directly counteracts the diverging effects of drift (García-Navas et al. 2015; Poelstra et al. 2014). Another possibility is that sufficient divergent selection could contribute to overcome the homogenizing effects of gene flow (Coyne & Orr 2004; Feder et al. 2013; Nosil & Feder 2012; Wagner et al. 2012; Wolf & Ellegren 2017). In light of this view, the “genomic islands of speciation” model was proposed to illustrate that divergent selection can have effects in specific regions across the genome leading to divergence with gene flow (Feder et al. 2013; Nosil & Feder 2012; Wagner et al. 2012; Wolf & Ellegren 2017). Data from several organisms suggest that incipient speciation can be maintained with divergence at a small number genomic regions (Martin et al. 2013; Morales et al. 2017; Nadachowska-Brzyska et al. 2013; Towes et al. 2016 but see Curickshan et al. 2014, Burri 2017).

The geographical boundary between the two plovers was ambiguously defined in previous studies (Kennerley et al. 2008; Rheindt et al. 2011). Our results show that the discontinuity in genetic structuring (Figure 2c) and morphometric values (Figure 1b-d) between the two species is situated at coastline between Wenzhou and Fuzhou, indicating a contact zone (Figure 2c). For incipient species, individuals may show a clinal pattern of allele frequency and morphology at their contact and form a hybrid zone (Bastos□Silveira et al. 2012; Harrison 1986; Taylor et al. 2014). Our STRUCTURE results revealed no obvious signs of a hybrid zone (Figure 2d), and rather indicated sporadic migrants in the respective range of the other species. One possibility is that our sampling transect was not fine-grained enough to be able to discover the potential hybrid zone, which would thus be narrower than the 200 km distance between the two neighboring sampling sites. Another possibility for an apparent lack of a hybrid zone is assortative mating between the two plovers at the contact (Bearhop et al. 2005). The latter explanation is supported by the fact that populations of the two species close to this region were much more divergent in morphological traits than ones farther apart (Figure 1b-d). Reproductive character displacement between the two species (Brown & Wilson 1956; Robinson & Wilson 1994; Grant and Grant 2006) in the contact regions would constrain interbreeding (Rybinski et al. 2016; Winkelmann et al. 2014). Obviously, genetic data alone cannot demonstrate this and detailed experiments related to assortative mating in the contact zone are required.

### Taxonomic implications

This study offers several taxonomic implications for the *C. alexandrinus* complex in East Asia. Kennerley et al. (2008) recommended that the tropical breeding population, previously defined as the subspecies *dealbatus* warranted species status, based on differentiation in morphology, behavior and distribution from *C. alexandrinus*. Uncertainty arouse about the taxonomic status because *dealbatus* was not distinguished from *C. alexandrinus* in a first genetic evaluation (Rheindt et al. 2011). Our results provide further support for a pair of incipient species, indicating that *C. dealbatus* is the youngest species within the genus *Charadrius* (Barth et al. 2013). Although we did not apply explicit Bayesian species delimitation analyses (e.g. Yang & Rannala 2014), our results clearly demonstrate multiple diagnosable morphological characters and distinct ecological niches, which are likely key factors to maintain species limits. Beyond the detection of differences between these plovers, this study forms a basis for conservation considerations of *C. dealbatus* because of its restricted range and adaptation to subtropical climate, as well as a risk of population decline caused by habitat loss in the China coastline (Ma et al. 2014).

### Conclusions and perspectives

Resolving the balance between diverging selection and gene flow is of fundamental importance to understand speciation processes. Here, we show that *C. alexandrinus* and *C. dealbatus* represent a case of incipient species in which divergent selection associated with ecological differences likely works as efficient mechanism for the maintenance of divergence in the face of gene flow. While the full species status of *C. dealbatus* may be justified, it remains untested whether a strong level of reproductive isolation has been established between *C. alexandrinus* and *C. dealbatus*. Further, we have shown low genetic divergence between the two plovers, so it would be of particular importance to explore the patterns of divergence at the genome level and determine whether specific regions are related reproductive isolation and adaptation. In addition, the IM analysis presented here estimated average historical gene flow, but it would be interesting to evaluate more realistic (and complex) demographic models in order to estimate changes in *N*e and the span of isolation and secondary contact better. Genome-wide population genetic approaches promise the power for better resolved evolutionary history of the two *Charadrius* plovers but also open an avenue to characterize the genetic architecture associated with phenotypic trait divergence and local adaptation (Freedman et al. 2014; Nadachowska-Brzyska et al. 2013; Towes et al. 2016).

## ACKNOWLEDEMENTS

We thank Qiaoyi Liang, Xiaoyan Long, Derong Meng, Demeng Jiang, Li Tian and Hebo Peng help with field sampling, Guoling Chen, Yangkai Zhou for their assistance with laboratory works. Special thanks are due Shaochong Peng for preparing sketches of plovers in Figure 1.

## DATA ACCESSIBILITY

Sequences deposited at GenBank: accession number xxxxx-xxxxx

Phenotype, distribution, stable isotope and microsatellite genotypes available at: Dryad Doi: xxx

## AUTHOR CONTRIBUTIONS

Y.L., Z.W.Z. and T. S. conceived and designed the study, P.J.Q., C-Y.C., Q.H., J.M., N.Z., X.J.W., and T. S. collected data in the field, X.J.W. and S.M.L conducted molecular genetic laboratory work, X.J.W. P.J.Q. analyzed genetic data with input from Y.L. and G.H.. J.H.H. performed niche modeling analysis, C.C.Z. carried out stable isotope analysis with input from E.P.-G. and C.D.. X.J.W P.J.Q. and Y.L. wrote the manuscript with input from J.H.H., G.H., X.C.Z. and C.D. All authors gave final approval for publication.

